# Aniridia-related Keratopathy Relevant Cell Signaling Pathways in Human Fetal Corneas

**DOI:** 10.1101/2021.03.19.436141

**Authors:** André Vicente, Marta Sloniecka, Berit Byström, Jing-Xia Liu, Fátima Pedrosa Domellöf

## Abstract

**Background:** To study aniridia-related keratopathy (ARK) relevant cell signaling pathways (Notch1, Wnt/β-catenin, Sonic hedgehog (SHH) and mTOR) in normal human fetal corneas in comparison with normal human adult corneas.

**Results:** 20 wg fetal and normal adult corneas showed similar staining patterns for Notch1, however 10-11 wg fetal corneas showed increased presence of Notch1. Numb and Dlk1 had an enhanced presence in the fetal corneas compared to the adult corneas. Fetal corneas showed stronger immunolabeling with antibodies against β-catenin, Wnt5a and Wnt7a, Gli1, Hes1, p-rpS6, and mTOR when compared to the adult corneas. Gene expression of Notch1, Wnt5A, Wnt7A, β-catenin, Hes1, mTOR and rps6 was higher in the 9-12 wg fetal corneas when compared to adult corneas.

**Conclusions:** The cell signaling pathway differences found between human fetal and adult corneas were similar to those previously found in ARK corneas with the exception of Notch1. Analogous profiles of cell signaling pathway activation between human fetal corneas and ARK corneas suggests that there is a less differentiated host milieu in ARK.

## 1 Introduction

Aniridia is a congenital autosomal dominant disease caused by haploinsufficiency of the PAX6 gene transcription factor.^1^ It includes, among other clinical features, aniridia-related keratopathy (ARK), which classically occurs after childhood and recurs even if the patients are submitted to corneal transplantation.^2, 3^ This chronic progressive keratopathy is characterized by a multitude of defects such as disturbed corneal limbal cell differentiation, fragile epithelial cells, compromised epithelial cell adhesion and a chronic wound-healing state with compromised barrier function, which result in epithelial erosions, corneal conjunctivalization and vascular pannus with significant impact on vision.^4–6^, We have recently reported altered cell signaling pathways in ARK, with decreased detection of the Notch1 cell signaling pathway and enhanced activation of the Sonic hedgehog (SHH), mTOR and Wnt/β-catenin cell signaling pathways not only in the epithelium but also in the subepithelial pannus.^7^ Given that many tissues recapitulate patterns of fetal maturation during regeneration, here we investigated whether these differences mimic the signaling pathway patterns present during human fetal corneal development, which may reflect the presence of a likewise immature host milieu in ARK corneas.

The development of the cornea includes two waves of neural crest cells and the primitive corneal epithelium appears around day 33. The epithelium is then two-layered and already has a basement membrane.^8^ At 7 weeks of gestation (wg), mesenchymal cells migrate to form the corneal stroma and endothelium and at 7.5 wg the keratoblasts, which will later differentiate into keratocytes, are arranged in 4-5 incomplete layers with few collagen fibrils. The corneal epithelium continues to evolve into a stratified squamous epithelium^9^ and by 11-12 wg all corneal layers are formed, with the exception of the Bowman’s layer.^8^ The epithelium has then 2-3 cell layers and the stroma has 25-30 layers of keratoblasts and is separated from the endothelium by an irregular Descemet’s membrane.^10^ The Bowman’s layer is formed around 16-17 wg^11^ and by then the keratoblasts are distributed in a disorganized pattern in the anterior stroma.^10^ At 7 months of gestation, the cornea already has an adult structure but in the anterior stroma there are more keratoblasts and the collagen lamellae are still more randomly oriented.^8^ The corneal structures generally develop adult features after the first two years of life.^9^, ^12^

Notch1, Wnt/β-catenin, SHH, and mTOR cell signaling pathways are essential for eye development, regulate both cell proliferation and homeostasis in mammals and are altered in ARK corneas.^7,13–16^ The Notch1 signaling pathway coordinates corneal epithelial repair, vertical migration and regulation of basal corneal epithelial cells.^35^ In the central corneal epithelium, these processes depend on transient amplifying cells (TAC), which have high migratory and proliferative capacity.^17^ TAC are derived from limbal epithelial stem cells during normal corneal epithelial homeostasis and wound healing processes and migrate centripetally in the basal layer to maintain the corneal epithelium.^18^ Dlk1, a transmembrane protein that functions as negative regulator of Notch1, ensures that cells are kept in a progenitor mode, which is essential in developmental processes.^19^ We have previously shown that Dlk1 activation is increased in ARK corneal epithelial cells and stroma. Numb, another important negative regulator of Notch1, is also upregulated in ARK corneas.^7^

Determination of limbal stem cell fate is regulated by the Notch1 and Wnt/β-catenin signaling pathways and are essential during normal tissue development and regeneration.^16, 20^ The extracellular ligands Wnt5a and Wnt7a stimulate the Wnt/β-catenin signaling cascade, which leads to proliferation of corneal limbal stem cells^21^, and is increased in ARK corneas.^7^ β-catenin is important for the regulation of epithelial differentiation and stratification.^22^

The SHH pathway regulates maintenance of cell polarity, cell differentiation and proliferation and leads to activation of the glioma-associated oncogene homolog (Gli1) transcription factor that controls cell growth and survival.^23^ This pathway is upregulated in case of corneal epithelium debridement^24^ and in ARK corneas.^7^ It shares a downstream effector, Hes1, with the Notch 1 signaling pathway, which is essential for the regulation of corneal epithelial stem cells and mammalian eye development^25^. The serine/threonine kinase mammalian target of rapamycin (mTOR) signaling pathway regulates cell growth and proliferation. Activation of the mTOR signaling pathway leads to phosphorylation of the ribosomal protein S6 (rpS6).^26^ mTOR signaling is upregulated in ARK corneas^7^ and premature upregulation of mTORC1 during development in mice causes aniridia and anterior segment dysgenesis.^15 15, 16, 27^

How these signaling pathways contribute to development of the human fetal cornea is currently poorly understood. In the present study we employed immunostaining in conjunction with fluorescence microscopy and qPCR to comparatively analyze potential divergence of the important developmental signaling pathways Notch1, Wnt/β-catenin, SHH and mTOR between fetal and adult human corneas.

Given that many tissues recapitulate patterns of fetal maturation during regeneration, here we investigated whether these differences mimic the signaling pathway patterns present during human fetal cornea development, which may reflect the presence of a likewise immature host milieu in ARK corneas.

## 2 Results

Although the epithelium was not optimally preserved in the fetal corneas, it was still possible to evaluate the staining patterns in all samples. The fetal corneas from 10-11 wg and 20 wg and the adult corneas presented similar patterns of immunolabeling with the antibody against Notch1, seen as streaks in the stroma and stronger staining around the basal layers of the corneal epithelium; however the 10-11 wg fetal cornea showed increased presence of Notch1 (fig. 1A-C). Dlk1 immunolabeling could be observed in the epithelial cells at both 10-11 wg and 20 wg. Numb labelling was observed as streaks in the stroma of all fetal corneas. Numb was more abundant in the anterior stroma (fig. 1D-E, asterisk). Conversely, Numb could not be detected in the stroma of adult corneas but was present in the epithelial cells (fig. 1F).

**Figure 1:**
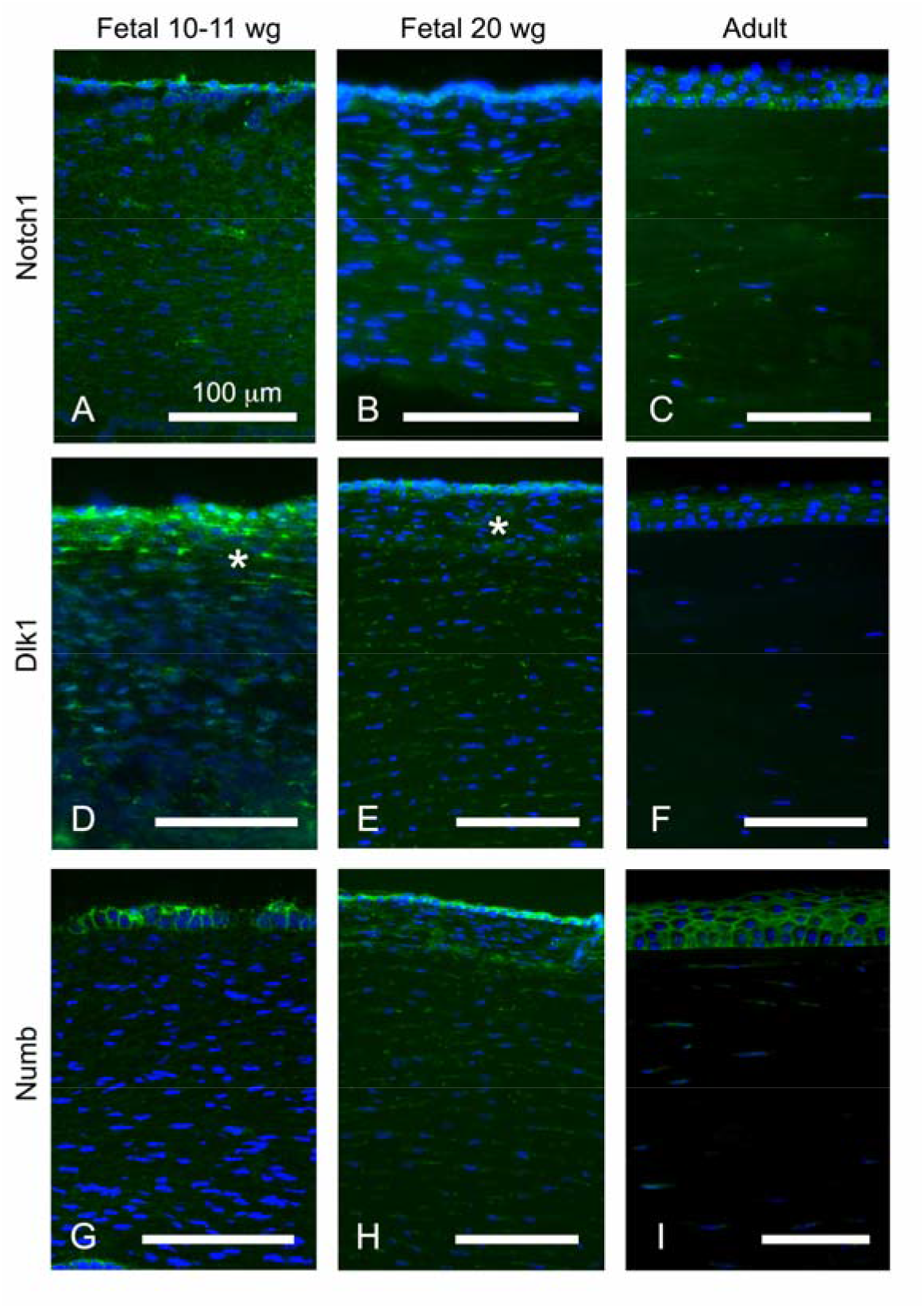
Cross-sections of fetal corneas 10-11 wg (A, D, G) and 20 wg (B, E, H) as well as normal adult corneas (C, F, I) labeled with antibodies (green) against Notch1 (A-C), Dlk1 (D-F) and Numb (G-I). The corneal epithelium is shown at the top and the stroma below, in all photographs of all figures. Cell nuclei are labeled blue with DAPI (A-I). Immunolabeling against Notch1 (A-C) was detected as streaks in the stroma and strongly around the basal layers of the epithelium in all fetal and adult corneas (A-C). In the 20 wg fetal corneas the labelling of the epithelium against Notch1 was slightly surpassed by the strong DAPI labelling of the epithelial cell nuclei (B). Dlk1 (D-F) labelled the epithelial cells and streaks in the stroma of all fetal corneas, more abundantly in the anterior region (D-E, asterisk), whereas in the adult corneas labeling was only present in the epithelium (F). In all fetal corneas, Abs against Numb (G-I), another inhibitor of Notch1, labelled the epithelial cells and streaks in the stroma (G-H) but stromal labelling was more abundant in the 20 wg (H) than in the 10-11 wg fetal corneas (G). Numb labeling in adult corneas was present at the epithelial cells and in sporadic streaks in the stroma (I). Bars: 100 μm.

Strong labeling with the Abs against Numb was observed in the epithelial cells of all fetal corneas, clearly delineating their contours, and in stromal streaks (fig. 1G-H). In the 20 wg fetal corneas (fig. 1H) these Abs labelled the stroma in streaks more abundantly than in the 10-11 wg fetal corneas (fig. 1G). In contrast, in the adult corneas, Abs against Numb labelled the epithelial cells but only sporadic streaks were present in the stroma (fig. 1I).

The epithelial cells of all fetal corneas were labelled with the Abs against Wnt5a (fig. 2A-B) and the stroma was immunolabeled with these Abs in streaks, more abundantly in the 10-11 wg (fig. 2A) than in the 20 wg fetal corneas (fig. 2B). In the 20 wg fetal corneas the immunolabeled streaks were slightly more abundant in the anterior stroma (fig. 2B). In contrast, only the epithelium was labelled with the Abs against Wnt5a in the adult corneas and the stroma was unlabeled (fig. 2C). Immunolabeling with Abs against Wnt7a was identified in the epithelial cells and as streaks in the stroma of all fetal corneas (fig. 2D-E). Labelling in the stroma of all fetal corneas presented as streaks which, as for Wnt5a, were slightly more abundant in the 10-11wg (fig. 2D) than in 20 wg corneas (fig. 2E). The Abs against Wnt7a labeled the epithelium in adult corneas but in the stroma only extremely sparse streaks were labelled (fig. 2F). The Ab against β-catenin intensely labelled the contours of the epithelial cells in a similar pattern in both the 10-11 (fig. 2G) and 20 wg (fig. 2H) fetal corneas. In the stroma of all fetal corneas, this Ab only labeled discrete streaks (fig. 2G-H). In the adult corneas, labeling with the Ab against β-catenin was completely absent in the stroma, but the epithelial cells were labeled in a pattern that was stronger in the basal region (fig. 2I).

**Figure 2:**
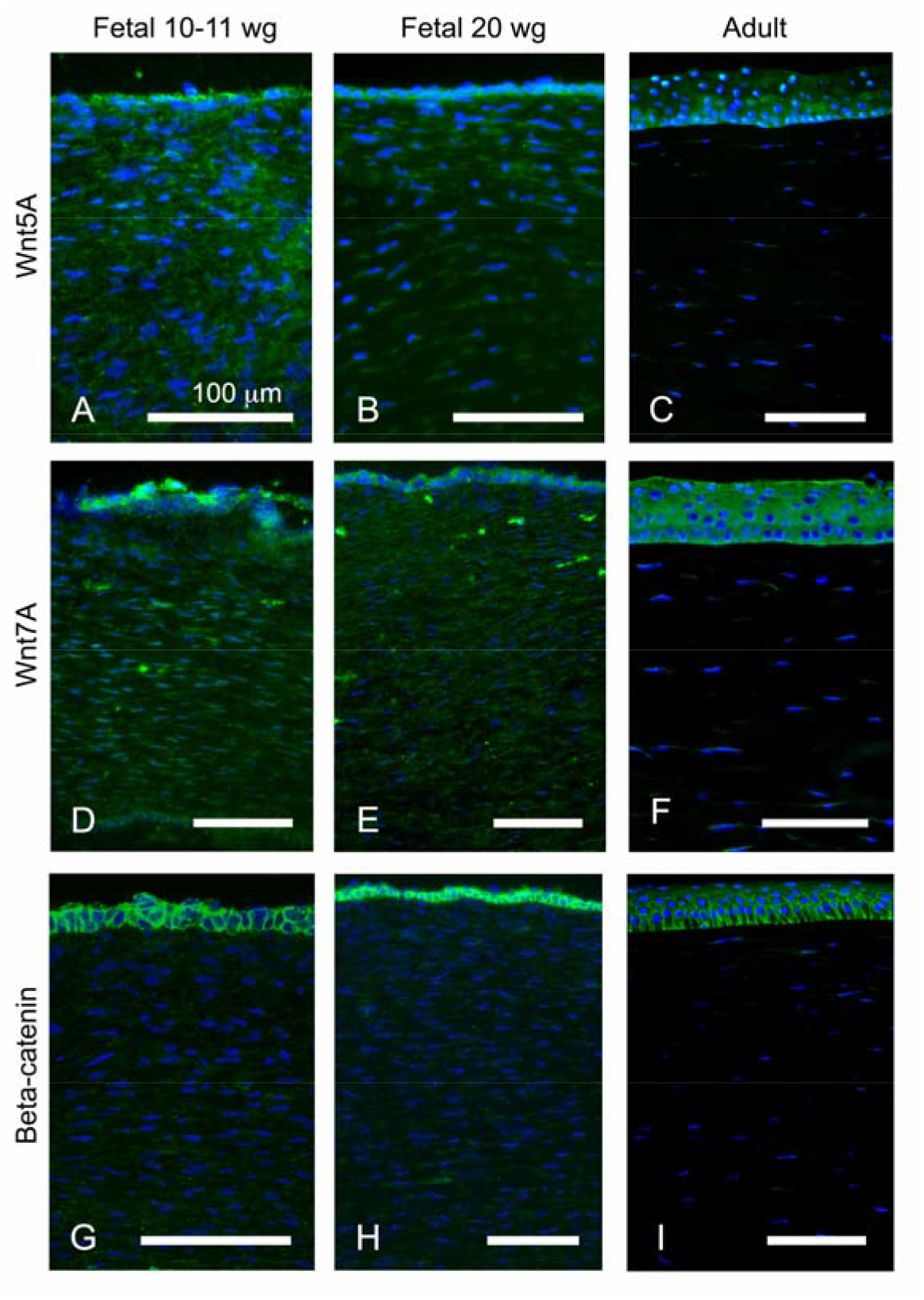
Cross-sections of fetal corneas 10-11 wg (A, D, G), 20 wg (B, E, H) and normal adult corneas (C, F, I) labeled with antibodies (green) against Wnt5a (A-C), Wnt7a (D-F), β-catenin (G-I). Cell nuclei are labeled blue with DAPI (A-I). The Abs against Wnt5a abundantly labelled both the epithelial cells and streaks in the stroma of the fetal corneas (A-B). The stromal labelling was more abundant in the 10-11 wg (A) than in the 20 wg fetal corneas (B), in which the streaks were more profuse in the anterior region (B). In the adult corneas, only the epithelium was labelled (C). Immunolabeling against Wnt7a was found in the epithelial cells of all fetal samples (D-E). The stromal labeling in all fetal corneas presented streaks, slightly more abundant in the 10-11 wg (D) than in the 20 wg fetal corneas (E). In contrast, in the adult corneas, labeling was present in the epithelium but only in extremely sparse steaks in the stroma (F). β-catenin immunolabeling was present abundantly in the contours of epithelial cells but only discretely in stromal streaks, in a similar pattern in the 10-11 wg (G) and 20 wg fetal corneas (H). In the adult corneas, labeling was present in the epithelial cells more intensively in the basal region, but absent in the stroma (I). Bars: 100 μm.

The Abs against Hes1 strongly labelled the epithelium of all fetal corneas (fig. 3A-B). These Abs labelled the stroma in streaks, more abundantly in the 10-11 wg (fig. 3A) than in the 20 wg fetal corneas (fig. 3B) and in the latter immunolabelling was more marked in the anterior stroma (fig. 3B). These Abs did not label the epithelium or the stroma in the adult corneas (fig. 3C). The epithelial cells in all fetal corneas were labeled with the Abs against Gli1 (fig. 3D-E) and the labelling was more abundant in the basal region of the epithelium in the 20 wg fetal corneas (fig. 3E). Immunolabelling with these Abs was found in streaks in the stroma of all fetal corneas (fig. 3D-E) and was stronger at 20 wg (fig. 3E), whereas it was completely absent in the epithelium and stroma of adult corneas (fig. 3F).

**Figure 3:**
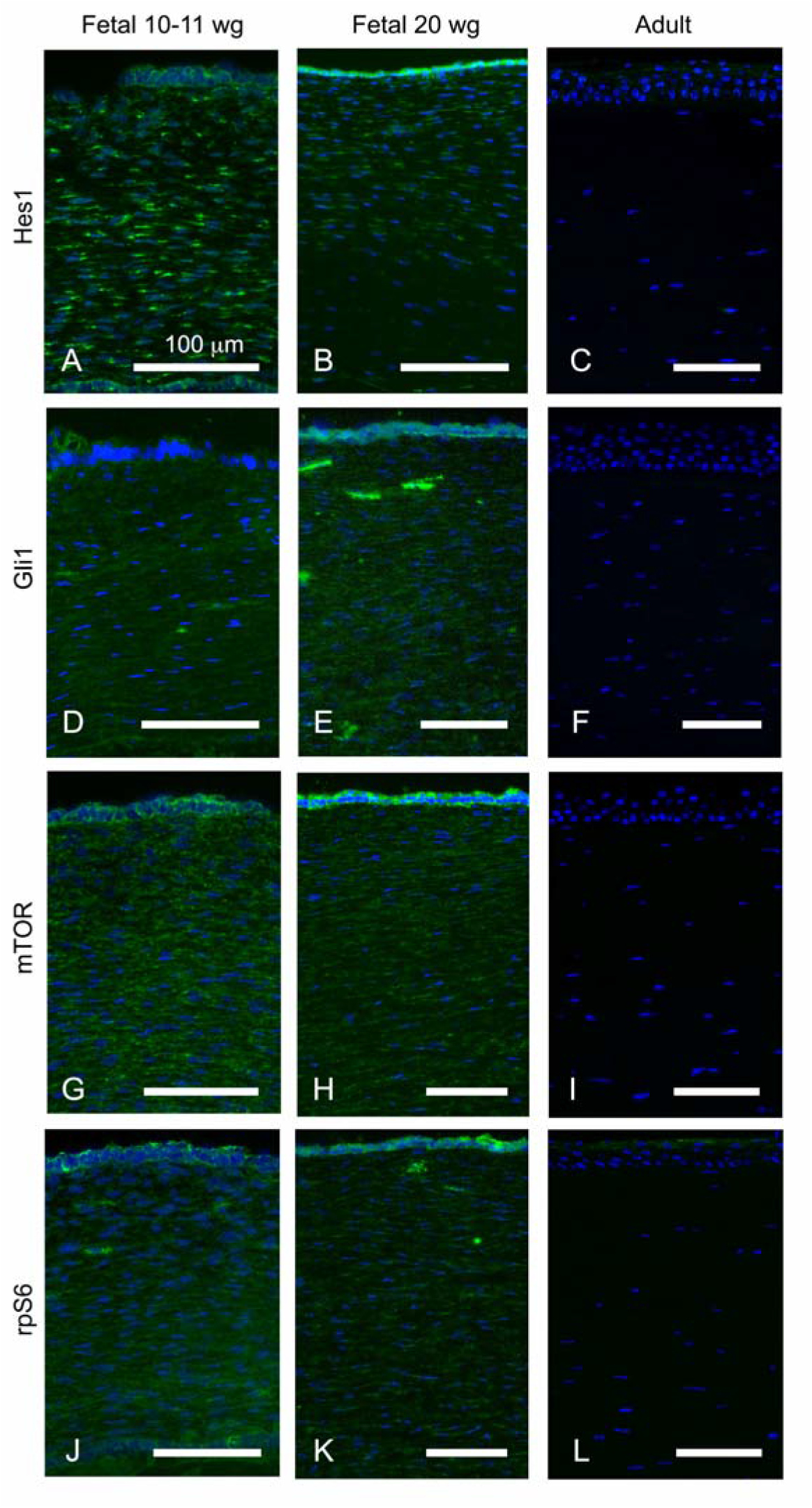
Cross-sections of fetal corneas 10-11 wg (A, D, G, J) and 20 wg (B, E, H, K) and adult corneas (C, F, I, L) labeled with antibodies (green) against Hes1 (A-C), Gli1 (D-F), mTOR (G-I) and p-rpS6 (J-L). Cell nuclei are labeled blue with DAPI (A-L). Abs against Hes1 (A-C) strongly immunolabeled the epithelial cells of the fetal corneas. These Abs labelled streaks in the stroma more profusely in the 10-11 wg (A) than in the 20 wg fetal corneas (B) and more intensively in the anterior stroma (B). In contrast, in the adult corneas, immunolabelling was not detected (C). Immunolabeling with Abs against Gli1was present in the epithelial cells in all fetal corneas (D-E), and was more intense in the basal region of the epithelium of the 20 wg fetal corneas (E). Labelling in the stroma was present in streaks in all fetal corneas (D-E) but was more abundant in the 20 wg fetal corneas (E). Immunolabelling for these Abs was not observed in adult corneas (F). The Ab against mTOR abundantly labeled the epithelial cells of all fetal corneas (G-H) but marked the epithelium in the 20 wg fetal corneas more intensively (H). The stroma in all fetal corneas was labelled in streaks (G-H) but was more abundantly marked in the 10-11 wg fetal corneas (G). This Ab did not immunolabel the adult corneas (I). The Ab against p-rpS6 labeled the epithelial cells and abundant streaks in the stroma of all fetal corneas in a likewise pattern in both 10-11 wg (J) and 20 wg fetal corneas (K). The stroma in the adult corneas was not labeled and the surface of the epithelium was only scarcely labelled, suggesting sticky adherence to the epithelial surface (L). Bars: 100 μm.

The Ab against mTOR labelled the epithelial cells in all fetal corneas (fig. 3G-H) but more strongly in the 20 wg fetal corneas (fig. 3H). The stroma of all fetal corneas presented labelled streaks (fig. 3G-H), which were more abundant in the 10-11 wg fetal corneas (fig. 3G), whereas in the adult corneas no immunolabeling against mTOR was detected (fig. 3I). Immunostaining against p-rpS6 was present in the epithelial cells and in abundant streaks in the stroma of all fetal corneas, with similar intensity in both 10-11 wg and 20 wg (fig. 3J-K). In the adult corneas, labelling with the Ab against p-rpS6 was absent in the stroma and it was only barely present in the surface of the epithelium, which could be indicative of non-specific adherence to the epithelial surface (fig. 3L).

We subsequently examined the gene expression profiles of the cell signaling pathways by RT-qPCR. The central part of the 9 wg, 10-11 wg fetal cornea, 12 wg fetal cornea, and 3 adult corneas were collected from frozen tissue sections using laser microdissection microscopy. In comparison to the adult corneas, expression of Notch1 gene was 3.5 fold higher in 9-12 wg fetal corneas (fig. 4A). Expression of Wnt5A, Wnt7A and β-catenin genes was increased by 1.7, 3.75 and 2.14 fold, respectively (fig. 4B). Gene expression of Hes1, mTOR and rps6 was increased by 2.2, 2.5 and 1.74 fold respectively (fig. 4C).

**Figure 4:**
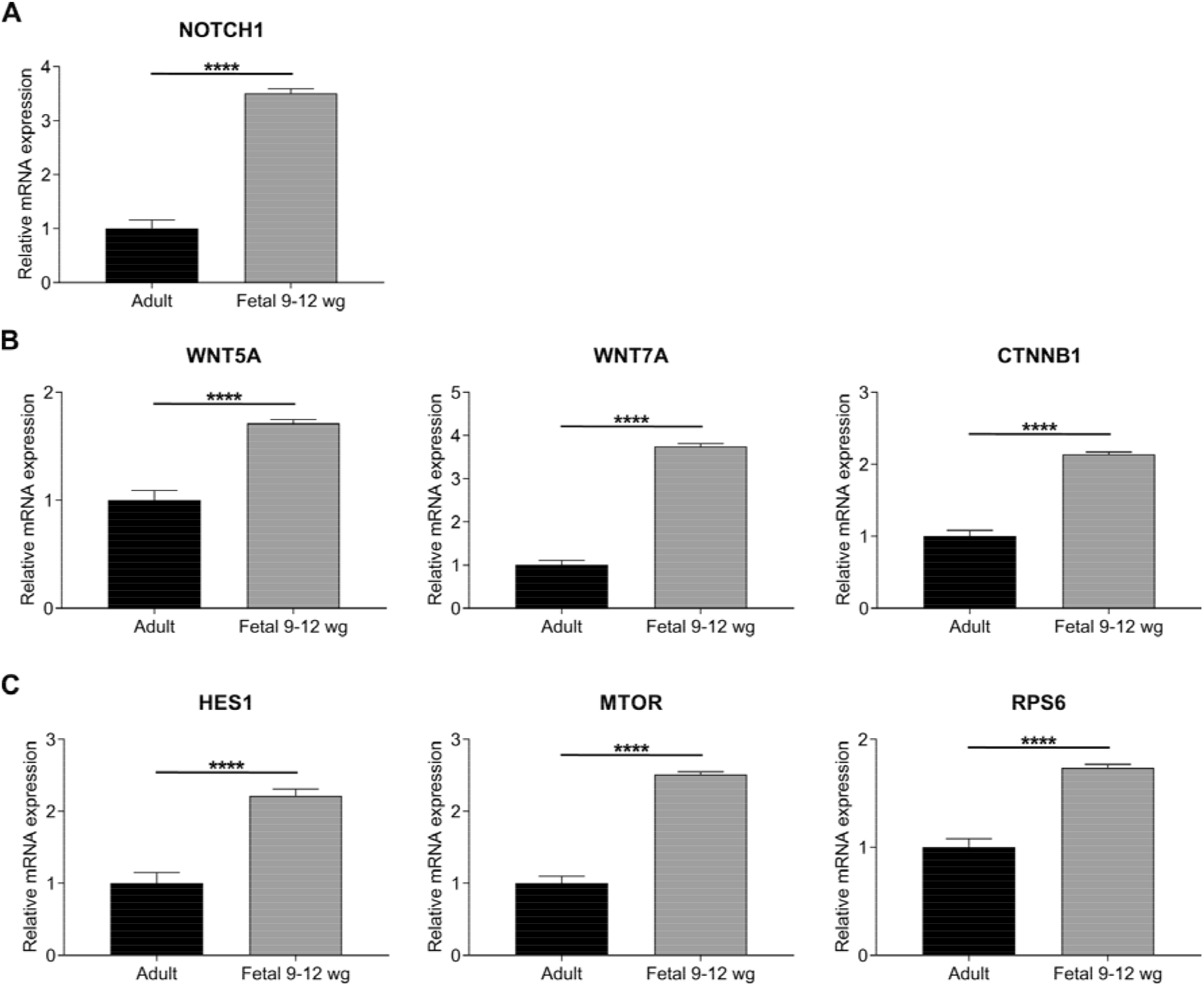
Gene expression of Notch1 (NOTCH1) (A), Wnt5A (WNT5A), Wnt7A (WNT7A), β-catenin (CTNNB1) (B), and Hes1 (HES1), mTOR (mTOR), rps6 (RPS6) (C) in 9-12 wg fetal corneas as compared to adult cornea. 3.5 fold increase in Notch1 gene expression was detected in 9-12 wg fetal corneas. (A). Expression of Wnt5A, Wnt7A and β-catenin genes was increased by 1.7, 3.75 and 2.14 fold, respectively. (B). Gene expression of Hes1, mTOR and rps6 was increased by 2.2, 2.5 and 1.74 fold respectively (C). Values are means ± SD. ****p<0.0001.

## 3 Discussion

The present study is the first to investigate the patterns of immunolabeling and gene expression of key cell signaling pathways related to cell proliferation and differentiation (Notch1, Wnt/β-catenin, SHH and mTOR) in the human fetal cornea. Cell differentiation is altered in ARK^5, 6^ and we have previously described changes in these cell signaling pathways in naïve aniridia corneas and in failed corneal transplants with advanced ARK.^7^ The present study allows a comparison between human fetal, normal adult and ARK corneas addressing the question whether there is an immature host milieu in ARK that mimics that encountered during fetal maturation.

The main limitation of this study is the reduced number of human corneal fetal samples derived from the scarcity of opportunity to collect such samples. On the other hand, the fact that the data was collected on human samples is a major strength of this study. We have previously reported reduced immunolabeling detection of Notch1 and increased detection of negative regulators of this signaling pathway (Dlk1 and Numb), in the epithelium and stromal pannus of ARK corneas^7^ when compared to normal adult corneas. Notch1 is responsible for corneal epithelial cell differentiation^28^ and lack of Notch1 in a mouse model leads to keratinized hyperplastic, skin-like corneal epithelium with neovascularization in the stroma,^29^ which reproduces, in part, the changes encountered in ARK corneas.^3^ In contrast, in fetal corneas, immunolabeling with Notch1 was detected in a similar pattern to that of normal adult corneas with 10-11 wg fetal corneas showing slightly increased presence of Notch1. Notch1 gene expression analysis revealed that it is expressed more abundantly in the fetal corneas (fig 4A). Nevertheless, the negative regulators of Notch1, Dlk1 and Numb, were also present in the stroma of the fetal corneas, as assessed by immunolabeling, but in the adult corneas they were only present in the epithelial cells. In the stroma of adult corneas, Dlk1 was absent and Numb was only present in extremely sporadic streaks in a pattern that was opposite to that observed in fetal and ARK corneas. This cell signaling pathway has a very diverse role in development and in self-renewing tissues, such as the corneal epithelium, having functions that range from serving as a gatekeeper for progenitor cells, cell lineage differentiation and barrier function regulation^28^ and we have now described a stronger detection of the key components of the Notch1 pathway in human fetal corneas during development. Dlk1 helps cells remain immature and secures progenitor cell populations^30^ and is important in fetal development. Here, we have revealed that it was abundantly expressed in the epithelial cells and stroma of 10-11 wg and 20 wg fetal corneas (fig. 1D-F) in a similar pattern to that previously reported in ARK corneas.^7^ Numb, also a Notch1 negative regulator presented a similar pattern of distribution to Dlk1 in the fetal corneas (fig 1G-I). There is a lack of data on the possible roles of Numb and Dlk1 in the corneal stroma. Numb has been shown to contribute to the maintenance of an undifferentiated state in epithelial progenitor cells in epidermis^31^ and the present findings can be interpreted as indicating the possibility of a similar role in the human cornea.

The Wnt/β-catenin cell signaling pathway has a decisive role in determining cell fate, proliferation, differentiation and apoptosis during development, as well as maintenance of stem cells and homeostasis in adult tissues^32^ and its presence is enhanced in the epithelium and pannus of corneas with advanced ARK.^7^ In the present study we showed that in fetal corneas at 10-11 wg and 20 wg, Wnt5a, and Wnt7a were abundantly identified in the corneal epithelial cells and stroma, whereas their presence was practically limited to the epithelium in normal adult corneas, as assessed by immunostaining. Upregulation of this signaling pathway in fetal corneas (as shown by both immunolabelling and gene expression analysis), was similar to what has been described in other ocular tissues during eye development.^33, 34^ A previous study examined the importance of the Wnt/β-catenin cell signaling pathway in corneal development in mice and established that conditional ablation of β-catenin led to precocious stratification of the corneal epithelium, postulating that this signaling pathway plays a crucial role in epithelial stratification in corneal morphogenesis.^22^ Furthermore, it has been shown that the Wnt/β-catenin cell signaling pathway influences the development of different ocular cell types in fetal corneas, including epithelial limbal stem cells, epithelial cells^35,36^ and keratocytes. In addition, it has been demonstrated before, in a mouse model, that Wnt/β-catenin cell signaling is not present in adult corneal limbus but that it is active in wing and squamous corneal epithelium and in developing corneal stroma and endothelium, in contrast to what happens in adult stroma and endothelium.^16^ Here we have shown that the staining patterns regarding the Wnt/β-catenin signaling pathway in the epithelium and stroma of human fetal corneas seem to mimic a pattern of activation of this signaling pathway described previously in mice.^22^ This signaling pathway seems to have a complex and multifaceted role in the development and maintenance of the corneal epithelial cells.^16^

SHH is a secreted protein that works as a morphogen during development^37^ and directly regulates the cell cycle, promoting proliferation by upregulating cyclin D1.^38^ Here we found a stronger detection of elements of the SHH signaling pathway (Gli1 and Hes1) in 10-11 wg and 20 wg corneas, as well as increased Hes1 gene expression in 9-12 wg fetal corneas, suggesting that activation of this pathway might be important during fetal corneal development. Hes1, in particular, has a role in preventing differentiation and maintaining progenitor cells and proliferation during development.^25^ Its abundant detection in the epithelial cells and stroma of fetal corneas, compared with its absence in adult corneas, suggests a role in cell proliferation during development and mimics the pattern described in ARK corneas.^7^ SHH is upregulated in migrating corneal epithelial cells.^24^ Human mutations in SHH result in holoprosencephaly which includes anophthalmia and cyclopia.^39, 40^ Furthermore, it has been proven that loss of SHH signaling in the lens disturbs the migration of neural crest cells into the cornea.^41^ Therefore, SHH signaling has both direct and indirect effects on corneal development.^41^ The results in the present study further emphasized its importance during corneal fetal development and the important parallel with the patterns encountered in ARK corneas.^7^

The mTOR cell signaling pathway can be described as a central regulator of cell proliferation, growth, motility, transcription, protein synthesis, autophagy and survival.^15, 26^ It regulates apoptosis and cell proliferation in pterygium^42^ and was suggested to be upregulated in the corneal epithelium and pannus in advanced ARK.^7^ In the present study, both mTOR and p-rpS6, a downstream element of this cell signaling pathway, were found in the fetal corneal epithelial cells and stroma but were absent in normal adult corneas, suggesting that this signaling pathway is also upregulated in fetal corneas at 10-11 wg and 20 wg, where cell proliferation is most likely central. The gene expression analysis also showed that mTOR and rps6 expression are higher in fetal corneas. The contribution of this signaling pathway to proliferation has led to studies where its inhibition was used to treat corneal neovascularization^43^ and transplant rejection.^44^ The effects of rapamycin, an inhibitor of the mTOR signaling pathway, in human corneal epithelial cells *in vitro* have been previously tested, indicating that it prevents the loss of corneal epithelial stem cells to replicative senescence and apoptosis.^45^ Nevertheless, the effects of this signaling pathway on corneal development are not known and should be further explored.

Our findings related to the immunolabeling pattern for these cell signaling pathways in the basal layers of the corneal epithelium in fetal corneas, with labeling of areas usually populated by TAC in adults,^17^ raised attention to the importance of epithelial basal cells with high migratory and proliferative capacity in normal adults and ARK corneas. These cells are formed by amplification of epithelial limbal stem cells and migrate centripetally during normal corneal epithelial homeostasis and wound healing in response to growth factors, cytokines and changes in the extracellular matrix.^17, 18^ The importance of these processes and TAC in the chronic wound-healing in ARK remains a question to be further explored but there is an apparent similarity between the regulation of cell signaling pathways related to the activation of TAC in normal adults, epithelium in ARK corneas and in high proliferative fetal epithelial cells.

The PAX6 gene is known to regulate transduction in ocular cells and to have a crucial role in securing human corneal epithelium identity by regulating cell differentiation.^46^ We described previously how Notch1, Wnt/β-catenin, SHH and mTOR cell signaling pathways are altered in patients with ARK, a condition resulting from reduced Pax6 protein levels.^7^ With the exception of Notch1, the changes in the cell signaling pathways observed in the corneas of patients with ARK and the patterns detected on human fetal corneas at 9-12 wg, 10-11 and 20 wg were analogous. This suggests that in corneas with advanced ARK, with reduced Pax6 protein levels, the cellular microenvironment mimics a less differentiated milieu, similar to that occurring during normal fetal development in corneas with normal Pax6 protein levels.

In this study we analyzed localization and quantified the expression of Notch1, Wnt/β-catenin, SHH and mTOR in normal human fetal corneas and healthy adult corneas. Cell signaling in the human cornea during development is currently poorly understood but our data has highlighted that certain pathways including mTOR1 and SHH may be essential in differentiating epithelial cells. Similarity to what is present during normal corneal fetal development, a more undifferentiated milieu, supports the proposed importance of host-specific factors and the corneal microenvironment in the context of limbal stem cell deficiency in ARK. We found that there are substantial similarities, excluding activation of Notch1, between the gene expression profiles of key signaling components between fetal and ARK corneas.

## 4 Experimental Procedures

### 4.1 Corneal Samples

Eyes from eight human fetuses were collected, with ethical approval, after legal interruptions of pregnancy at 9 wg (n=2), 10-11 wg (n=2), 12 wg (n=2) and at 20 wg (n=2). Gestational age was calculated using the first day of the last menstrual period and confirmed with ultrasound. Five adult healthy corneas were collected from four male (74-, 76-, 82-, and 83-year-old) and one female (37-year-old) donors. These normal corneas were obtained from deceased individuals who, when alive, chose to donate their eyes post-mortem for transplantation and research purposes, according to Swedish law. The Regional Ethical Review Board in Umeå has determined the use of the post-mortem donated, anonymized tissue for study purpose to be exempt from the requirement for approval. The study followed the principles of the declaration of Helsinki.

The two human adult corneas (from 74- and 82-year-old males) used for immunochemistry were first fixed in formalin and embedded into paraffin wax. Superfrost^®^ Plus slides were used to collect serial sections, 5 μm thick, which were dried in a vertical position overnight at 60°C and then placed at + 4°C in a slide box, as previously described.^7^ The remaining three adult corneas and all fetal samples were mounted on cardboard with embedding medium (OCT cryomount, HistoLab Products AB, Gothenburg, Sweden), quickly frozen in propane cooled with liquid nitrogen and stored at −80°C. For immunohistochemistry 5 μm thick serial sections were cut at −23°C using a Leica CM3050 cryostat (Leica Microsystems, Wetzlar, Germany) and collected in Superfrost^®^ Plus slides (Thermo Fisher Scientific, Rockford, IL, USA). For laser microdissections, 10 μm thick serial sections of freshly frozen adult corneas, as well as 9 wg, 10-11 wg and 12 wg fetal corneas were cut in RNase free conditions at −23°C using a Leica CM3050 cryostat, and collected on NF1.0 PEN membrane slides (Zeiss, Jena, Germany). Sections were fixed in ice cold 70% ethanol for 1 minute and transferred to −80°C freezer until further processed.

### 4.2 Immunohistochemistry

The sections of the human fetal corneas were first brought to room temperature, left to dry (20 min) and then rinsed for three times (5 min) in 0,01M phosphate buffered saline (PBS). Sections from adult control healthy corneas were de-waxed with Tissue-Clear^®^ (1466, Sakura Finetek Europe, Alphen aan den Rijn, Netherlands) and rehydrated. Antigen retrieval was performed with a water bath at 95°C (30 min) in pre-warmed citrate buffer (10mM Citric acid, 0,05% Tween 20, pH 6,0) or Tris-EDTA buffer (10mM Tris Base, 1mM EDTA, 0,05% Tween 20, pH 9.0). The sections were put at room temperature (20 min) to cool down, washed in running water (5 min) and then submerged in 0,01M PBS, three times (5 min).

In order to block unspecific binding, fetal and adult corneal sections were incubated with 5% normal goat or donkey serum at room temperature (15 min). Thereafter, the sections were incubated with either monoclonal or polyclonal primary antibodies (Abs) specific against components of the Notch1 (Notch1; Dlk1; Numb), Wnt/β-catenin (Wnt5a; Wnt7a; β-catenin), Sonic Hedgehog (Gli1; Hes1) and mTOR (mTOR1; p-rpS6) signaling pathways (Table 1), at + 4°C overnight. The sections were subsequently incubated with the adequate secondary antibody (Table 2) at 37°C for 30 min, submerged in 0,01M PBS for three times (5 min) and then mounted with Vectashield^®^ mounting media with 4’,6-diamidino-2-phenylindole (DAPI; H-1000 and H-1200; Vector laboratories, Inc, Burlingame, USA). Potential unspecific binding was evaluated by omission of the primary antibody in control sections that were otherwise processed similarly.

**Table 1.** Primary Antibodies.

| Antigen | Antibody | Concentration | Source |
| --- | --- | --- | --- |
| Dlk1 | Ab21682 | 1:1000 | Abcam, Cambridge, UK |
| Notch1 | Ab52627 | 1:300 | Abcam, Cambridge, UK |
| Numb | Ab14140 | 1:250 | Abcam, Cambridge, UK |
| Wnt5a | Ab32572 | 1:100 | Abcam, Cambridge, UK |
| Wnt7a | Bs-6645R | 1:100 | Bioss Inc, Woburn, MA, USA |
| β-catenin | Ab32572 | 1:100 | Abcam, Cambridge, UK |
| Hes1 | GTX 108356 | 1:500 | Genetex, Irvine, CA, USA |
| Gli1 | Ab 92611 | 1:800 | Abcam, Cambridge, UK |
| mTOR | (7C10) 2983 | 1:500 | Cell Signaling, Danvers, MA, USA |
| p-rpS6 | GTX 60800 | 1:800 | Genetex, Irvine, CA, USA |

**Table 2.** Secondary Antibodies.

| Antigen | Antibody | Concentration | Source |
| --- | --- | --- | --- |
| Donkey anti-mouse DyLight 488 | 715-485-150 | 1:100 | Jackson ImmunoResearch Europe Ltd, Suffolk, UK |
| Goat anti-mouse Alexa 488 | A21121 | 1:300 | Molecular Probes, Life Technologies, Darmstadt, Germany |
| Donkey anti-rabbit FITC | 715-095-152 | 1:50 | Jackson ImmunoResearch Europe Ltd, Suffolk, UK |
| Goat anti-rabbit Alexa 488 | A11034 | 1:300 | Molecular Probes, Life Technologies, Darmstadt, Germany |

We have previously worked with both freshly frozen and formalin fixed corneal specimens, using the antibodies included in the present study. We have previously ascertained that the staining pattern was similar irrespective of the tissue being freshly frozen or paraffin embedded.^3^, ^12^, ^47^, ^48^

### 4.3 Image acquisition

The sections were photographed using a Leica DM 6000 B microscope (Leica Microsystems, Wetzlar, Germany), equipped with a digital high-speed fluorescence charge-coupled device camera (Leica DFC360 FX; Leica Microsystems, Wetzlar, Germany). Image processing was performed using Adobe Photoshop CS6 software (Adobe Systems, Inc., San Jose, CA, USA). Both the immunostaining procedures and image acquisition with fluorescence microscopy were performed using identical settings for all samples. Therefore, relative comparisons between different samples treated with the same antibody could be performed.

### 4.4 Laser microdissections

Before laser microdissection (LMD) was performed, the NF1.0 PEN membrane slides with 9 wg, 10-11 wg and 12 wg fetal corneas, and adult control corneas were transferred from −80°C directly to ice cold 70% ethanol for 2 minutes, washed in nuclease free water in order to remove residual OCT, and then dehydrated in 70, 96 and 100% ethanol for 20 seconds each. One additional wash in 100% ethanol was performed for 1 min. Afterwards, the slides were left to dry inside a fume hood for 10 minutes and transferred to a desiccator. LMD was performed using a PALM MicroBeam microscope (Carl Zeiss Microscopy, Jena, Germany). The central part of the corneas (stroma and epithelium) was carefully dissected, pooled (8 sections from each fetal cornea, and 16 sections from adult corneas), lysed in lysis buffer and used for gene expression analysis.

### 4.5 RT-qPCR

mRNA was extracted from the dissected corneas using RNA extraction kit (Qiagen, Venlo, Netherlands) according to the manufacturer’s instructions. Next, 35ng of RNA was reverse transcribed into cDNA using the high-capacity cDNA reverse transcription kit (Thermo Fisher). Notch1 (NOTCH1; Hs01062014_m1), Wnt5A (WNT5A; Hs00998537_m1), Wnt7A (WNT7A; Hs01114990_m1), β-catenin (CTNNB1; Hs00355045_m1), Hes1 (HES1; Hs00172878_m1), mTOR (mTOR, Hs00234508_m1), and rpS6 (RPS6; Hs01058685_g1) probes were used in order to determine gene expression (Thermo Fisher). Samples were run in duplicates in ViiA™ 7 Real-Time PCR System (Thermo Fisher). β-actin and 18S probes served as endogenous controls (Thermo Fisher). Analysis was performed with ViiA™ 7 Software (Thermo Fisher). The fold change gene expression in the fetal corneas at 9 wg, 10-11 wg and 12 wg was pooled together and compared to that of the adult corneas.

### 4.6 Statistical analysis

Data are presented as mean ± SD. Statistical analysis was performed with unpaired t-test. A p-value of < 0.05 was considered statistically significant.

